# Rapid, robust plasmid verification by *de novo* assembly of short sequencing reads

**DOI:** 10.1101/2020.07.02.185421

**Authors:** Jenna. E. Gallegos, Mark F. Rogers, Charlotte Cialek, Jean Peccoud

**Affiliations:** Department of Chemical & Biological Engineering, Colorado State University; GenoFAB, Inc; Department of Biochemistry, Colorado State University

**Author notes:** The authors wish it to be known that, in their opinion, the first two authors should be regarded as joint First Authors.

## Abstract

Plasmids are a foundational tool for basic and applied research across all subfields of biology. Increasingly, researchers in synthetic biology are relying on and developing massive libraries of plasmids as vectors for directed evolution, combinatorial gene circuit tests, and for CRISPR multiplexing. Verification of plasmid sequences following synthesis is a crucial quality control step that creates a bottleneck in plasmid fabrication workflows. Crucially, researchers often elect to forego the cumbersome verification step, potentially leading to reproducibility and— depending on the application—security issues. In order to facilitate plasmid verification to improve the quality and reproducibility of life science research, we developed a fast, simple, and open source pipeline for assembly and verification of plasmid sequences from Illumina reads. We demonstrate that our pipeline, which relies on *de novo* assembly, can also be used to detect contaminating sequences in plasmid samples. In addition to presenting our pipeline, we discuss the role for verification and quality control in the increasingly complex life science workflows ushered in by synthetic biology.

## Introduction

As synthetic biology programs increase in scale (1-3) workflows involving the high-throughput construction of dozens or even hundreds of plasmids are becoming increasingly common (4-6). DNA sequencing is an integral part of fabrication workflows involving the assembly of synthetic DNA fragments (7-10). Sequencing data can detect single point mutations (SNP) resulting from mistakes in the DNA synthesis processes (11-13) or from PCR. In addition, sequencing data can be used to detect structural issues such as inversion or duplication of genetic elements introduced during plasmid assembly.

Traditionally, plasmids have been verified using Sanger sequencing. This sequencing method requires a short, known sequence to initiate an up to ∼1000 nucleotide read. Typically, Sanger sequencing is used to verify a sequence of interest that has been inserted into a plasmid backbone by sequencing from known universal primer binding-sites on either side of the multiple cloning site, providing 2x coverage of the insert. Many bioinformatics applications used to edit plasmid sequences have features that facilitate the visual inspection of the alignment of the sequence reads and the plasmid sequence. This approach is practical to verify the insert of a limited number of plasmids. The quality and length of Sanger sequencing reads also simplify sequence assembly.

However, Sanger sequencing is not a viable quality control option for verifying sequences of a large number of plasmid libraries. In order to sequence whole plasmids by Sanger sequencing, the user must first design and order primers, which adds to the time and cost involved in verification. Depending on the sequence of the plasmid, it may be difficult to design primers that will generate ample coverage, and structural features like hairpins can result in low quality reads (14). In addition, the analysis of sequencing reads is time consuming and error prone.

In order to streamline this process, we previously developed an application to automate assembly and sequence verification of plasmid sequences from Sanger sequencing reads (15). However, because this pipeline relies on reference-based assembly, it struggles to resolve duplications, inversions, and other rearrangements that result in structural issues.

Recently, synthetic biologists have demonstrated the possibility of sequencing multiple plasmids in a single Oxford Nanopore run (16). This multiplexed approach overcomes many of the limitations of Sanger sequencing. Oxford Nanopore sequencing is fast, inexpensive, and does not require primers. However, the analysis of data still relies on a reference-based assembly.

Reference-based assembly is disadvantageous for quality control workflows because it is biased towards a particular goal. If there are contaminating sequences present in a sample, for instance, the reads for those contaminants may be thrown out during the assembly process. Reference-based assembly also requires a full reference sequence by definition. However, researchers may sometimes wish to identify unknown plasmid samples, and even for known samples, a full reference sequence is not always available. It is very common for plasmids described in the literature to be accompanied only by a visual map or vague descriptions of how the plasmid was constructed. Without a full reference sequence, it is not possible to design primers for Sanger sequencing or conduct a reference-based assembly.

To address these limitations, we present an alternative approach based on short read sequencing and *de novo* assembly. The costs associated with short read sequencing are rapidly decreasing while speed and read length are increasing (17, 18). The small amount of coverage needed for plasmids and various multiplexing options have also made short read sequencing a cost-effective option, as plasmid samples can easily be included within larger sequencing runs. Further, short read sequencing does not rely on primers and is less sensitive to secondary structures than long read sequencing. Though de novo pipelines exist for circular microbial genomes, these tools do not translate well to assembling plasmids, which are typically much smaller in size. DNA synthesis companies routinely rely on short read sequencing workflows for quality control in plasmid synthesis, but their assembly and verification tools are proprietary, and likely rely on reference-based assembly. Proprietary programs also exist in academic labs. However, our approach is the first open source, publicly available solution.

In this manuscript, we describe a plasmid verification pipeline that uses Illumina sequencing reads for *de novo* assembly. The pipeline was tested on a library of 96 plasmids designed to represent a broad range of variations of a common plasmid template. The tool produced correct assemblies for all 96 plasmid samples even when overall read quality was very low. Furthermore, we demonstrate that our script can determine whether or not a pool of reads is likely to contain contaminating sequences. We built a contamination test into our verification pipeline that informs the user of the likelihood of contamination. This tool can thus be used to verify known plasmid sequences, identify unknown plasmid sequences, and detect contamination in plasmid samples. The workflow described in this manuscript relies entirely on open source tools, making it, to our knowledge, the first non-proprietary tool for *de novo* assembly of plasmid sequences from short sequencing reads, and the first sequence assembly and verification tool that also predicts the likelihood of contamination.

## Materials and Methods

### Plasmid preparation and sequencing

We built and tested our pipeline using Illumina sequencing reads from 96 plasmid DNA samples. The sequencing dataset used to develop our workflow was generated in conjunction with a separate publication. Detailed explanations and plasmid reference sequences can be found in that manuscript (19).

Briefly, 6-24 individual transformants of eight different plasmids (ranging in size from 2521-3294 bp) were sequenced. Six of these plasmids are part of a family of synthetic plasmids that were synthesized, and sequence verified by Twist Biosciences (twistbioscience.com). Five of these six differ from each other by one to four SNPs which were intentionally introduced. These are referred to as “known-SNPs”. The sixth plasmid from Twist differs in that it lacks a 608bp insert present in the other five. This variant was used to analyze the impact of contaminants containing insertions and deletions (INDELs). The remaining two plasmids are unrelated and were generated by Gibson assembly (20) of parts that have not been sequence verified. These were, therefore, expected to contain a small and variable number of unknown SNPs and INDELs.

Plasmids were isolated from *E. coli* cells, analyzed on an Agilent TapeStation, and submitted to seqWell (seqwell.com) for sequencing. seqWell uses a library prep technology called plexWell that enables the preparation of hundreds or thousands of multiplexed samples for Illumina sequencing in just 3 hours.

Sequencing resulted in 2.9 million read pairs for the 96 samples (>1,000x coverage), with read lengths of 35 to 251 bases and mean per-read PHRED quality scores (21) ranging from 2.0 to 38.8 (average of 35.3). Optimal filtering conditions removed 93% of reads, yielding 1,682 to 11,938 filtered read pairs per plasmid with lengths of 125 to 251 bases and mean per-read quality from 36.8 to 38.8 (average of 38.0). In addition to raw reads in the form of FASTQ files, seqWell used their proprietary bioinformatics workflow to analyze the sequencing reads. They provided assembled FASTA files for 95 of the 96 samples.

### *De Novo* Assembly and Sequence Verification Pipeline

Our sequence verification pipeline performs three key steps: quality filtering, plasmid assembly and assembly evaluation. It accepts as input, Illumina paired-end sequencing FASTQ files containing sequencing reads and produces as output a FASTA assembly file representing the predicted plasmid reference sequence, along with an estimated likelihood of contamination (Figure 1). Reference sequences were used to build and evaluate the pipeline; however, it is important to emphasize that the pipeline itself does not use a reference.

**Figure 1.**
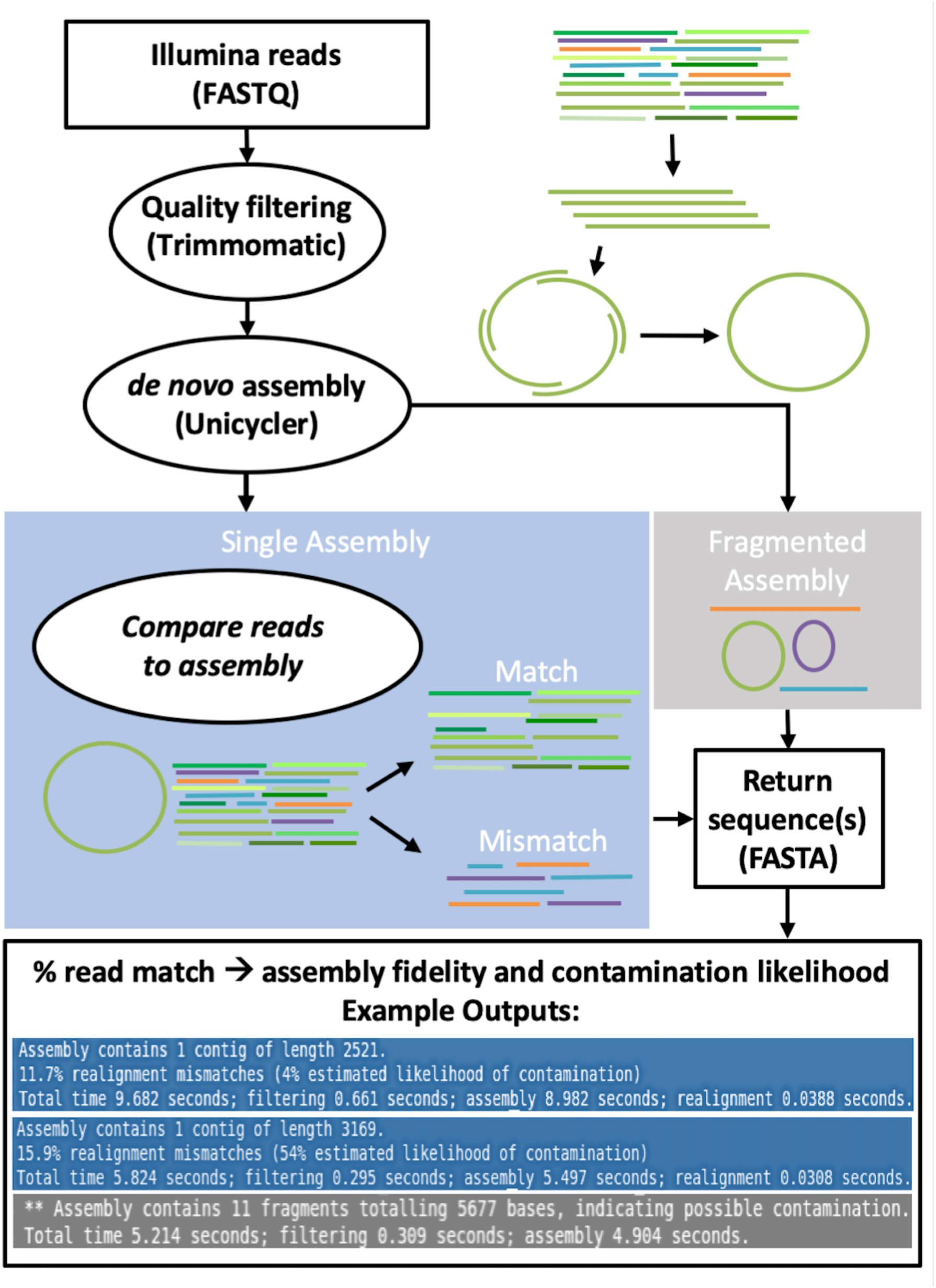
Schematic of pipeline. Illumina reads in the form of FASTQ files are filtered using Trimmomatic and assembled using Unicycler with parameters that have been specifically optimized for plasmid assembly. If a single assembly results, the unfiltered reads are compared with the assembled sequence and sorted according to whether or not they match 100%. Assembled sequences are returned as FASTA files. The percentage of matched reads is used to calculate the likelihood of contamination.

The first step is to filter the input sequences to eliminate all but the highest-quality reads. We tested two filtering tools, Trimmomatic, a commonly used open source software tool (22), and fastp, a more recent, high-throughput method, for comparison (23). Illumina sequencing can produce extremely high coverage for short plasmid sequences. Our experimental samples provided anywhere from 36,000 to 166,000 reads per plasmid, yielding coverage from 2,800x to over 12,000x. These values are orders of magnitude higher than typical recommendations for *de novo* genome assembly, which range from 13x to 60x (24, 25). Higher coverage does not improve performance, and often hurts performance, sometimes yielding assemblies with multiple sequences (putative contigs) (26). For this reason, we used stringent criteria for filtering reads, retaining only the top 3% to 11% highest quality reads.

The second step is sequence assembly, in which two FASTQ files containing trimmed, forward and reverse, Illumina paired-end sequencing reads are accepted as input, and a FASTA file containing one (or more) circular contig(s) is produced as output. For this, we considered five different assembly tools for our process: Unicycler (27), MIRA (28), Velvet (29), plasmidSPAdes (30) and Circlator (31). MIRA and Velvet are popular, general purpose assemblers that generate linear assemblies. Circlator is a post-processing tool that can convert linear assemblies to circular assemblies, while plasmidSPAdes and Unicycler are both designed to resolve chromosomes and plasmids from whole bacterial genome sequencing, yielding circular plasmid assemblies. We used MIRA and Velvet on their own and with Circlator, and we used Unicycler and plasmidSPAdes by themselves.

When assessing the quality of assembly of putative contigs, we imposed strict criteria: (1) the final assembly must contain only one contig that matches the correct plasmid sequence exactly, and (2) the pipeline should take no more than a few minutes to run.

### Assessing assembly quality and detecting contamination

Because our pipeline is intended for use in the absence of a reference sequence, once our pipeline was established, it was necessary to develop additional reference-agnostic methods for assessing plasmid assembly. For this, we built an additional quality control step into the pipeline in which the filtered reads are mapped back to the assembly. The proportion of reads that matches the assembled contig exactly is used to assess the quality of the assembly.

The proportion of reads that matches the assembled contig was further used to assess the likelihood of contamination in the original sample. There are several scenarios by which a plasmid sample might become contaminated. For instance, if plasmid DNA is extracted from a mixed population of transformants, depending on the method of plasmid construction and screening, there could be sequences present that have SNPs or that lack or contain an extra insert. Additionally, if a contamination event occurs between different laboratory strains, sequences from completely unrelated plasmids may be present in the mixed sample. In each case, when the contaminated sample is sequenced, the resulting reads will include a combination of reads from the primary sample and reads from the contaminant.

To simulate each of these possible contaminations (SNPs, INDELs, and unrelated sequences), we conducted a series of contamination simulation experiments by randomly sampling the filtered reads from distinct plasmids to create artificial contamination, and then built a procedure for detecting the contamination.

In each of these contamination simulation experiments, we select one of our plasmids and label it the “primary” plasmid, and its filtered reads, the “primary” read set. We then select a second, “contaminant” plasmid at random that has a different reference sequence from that of the primary plasmid. In each contamination simulation experiment, a subset of reads from the contaminant library is artificially combined with a subset of reads from the primary library, the reads are assembled, and the resulting assemblies are assessed. Different contamination levels (up to 50%) were assessed.

Our contamination detection algorithm uses simple string matching to compare an output assembly with the filtered reads used to create it. The algorithm iterates over all of the distinct high-quality reads, attempting to match them to a location in the assembly sequence. It assesses the percentage of distinct reads that map exactly to a location in the assembly and compares it to distributions we have established experimentally.

## Results

### Plasmid Identification and Verification

For quality filtering, we found that Trimmomatic (22) yielded reliably good data sets when we applied strict quality filtering that retained only the highest-quality 3% to 11% of read pairs (Figure 2). Users of our script may choose instead to use *fastp* (23) for quality filtering, since we found that both methods yielded similar results.

**Figure 2.**
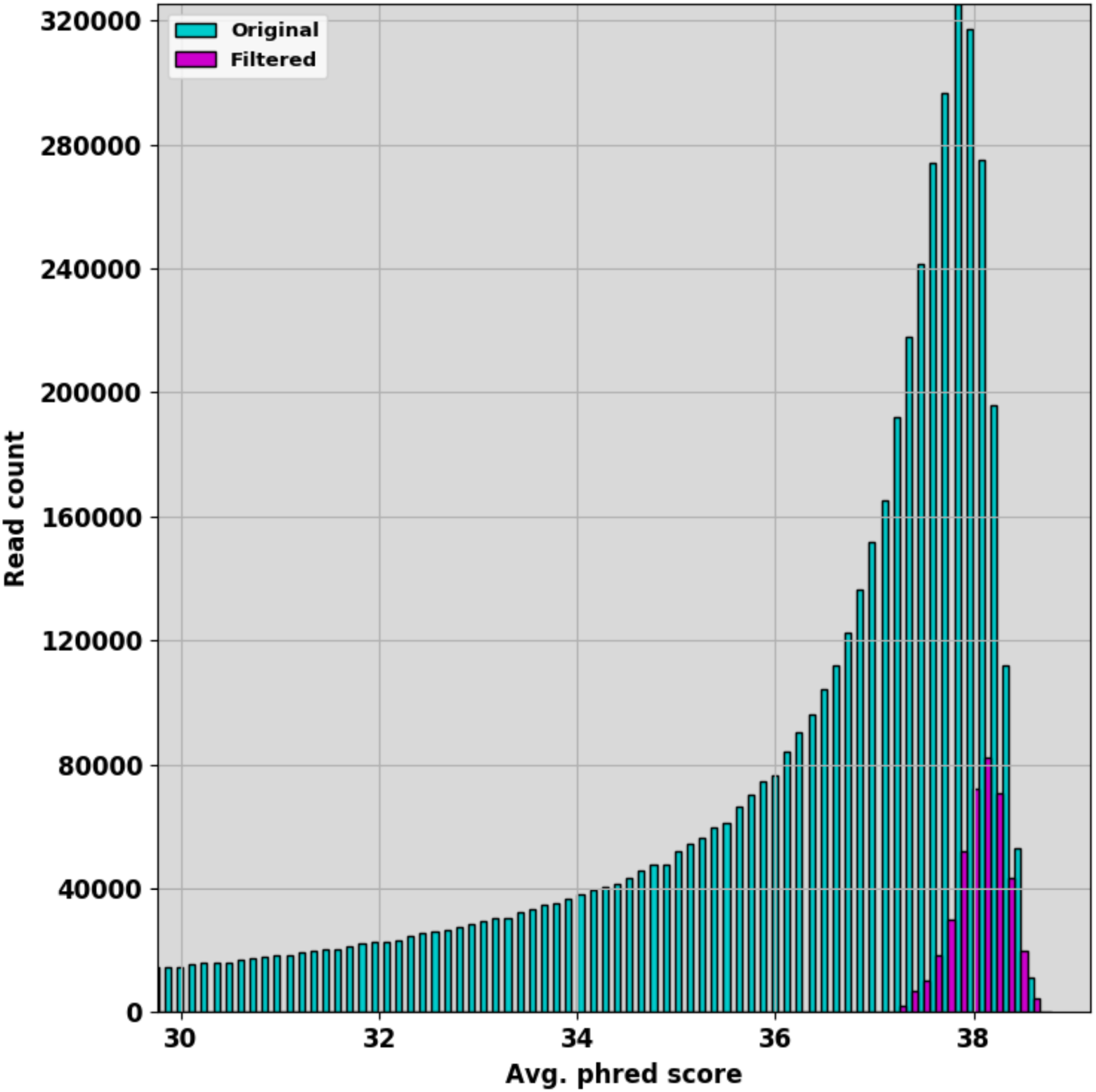
Filtering for high quality reads. Distributions of read qualities for original raw reads (cyan) and reads after strict filtering (7.1% of the original reads, magenta) show the dramatic increase in quality consistency in the reads provided to the assembler.

For assembly, the only tools that yielded single-contig assemblies in any of our tests were plasmidSPAdes and Unicycler. For our filtered data, plasmidSPAdes yielded 63/96 correct assemblies (66%) while Unicycler resolved all 96 correctly. Thus, we settled on the Unicycler program because we were able to derive quality assemblies with fewer parameter adjustments. It is worth noting that Unicycler is really a sophisticated wrapper for SPAdes (32).

Interestingly, none of the assembly tools we tried worked well with whole sets of Illumina reads even though the data for most of the samples were relatively high quality (average PHRED score of 35.3, Figure 2). The most likely explanation is an overabundance of input data, a paradox that poses a significant challenge for *de novo* assembly when sequencing coverage exceeds 500x (26).

Thus, we tested whether high-quality reads or the number of reads were necessary to produce accurate assemblies. We ran Unicycler on random samples of read pairs such that the resulting input data contained the same number of pairs as the quality-filtered data sets. We found that the randomly sampled read pairs yielded assemblies with the same fidelity as those built from high-quality reads (Supplemental Figure TBD or data not shown). However, we persisted in using quality filtering, as it reduced read quantity to a manageable level and the high-quality reads became important when predicting contamination, described below and in the next section. Using read filtering, optimal assemblies were achieved using coverage between 113x and 1030x (data not shown).

For plasmids ranging from 2.5 to 3.3kb, complete assemblies by Unicycler with our plasmid-specific parameters required on average less than eight seconds (single processor on a Linux laptop with 16GB RAM), including Trimmomatic filtering. The Unicycler assembly process itself dominated this time, which increased approximately linearly with the number of input reads (Figure 3).

**Figure 3.**
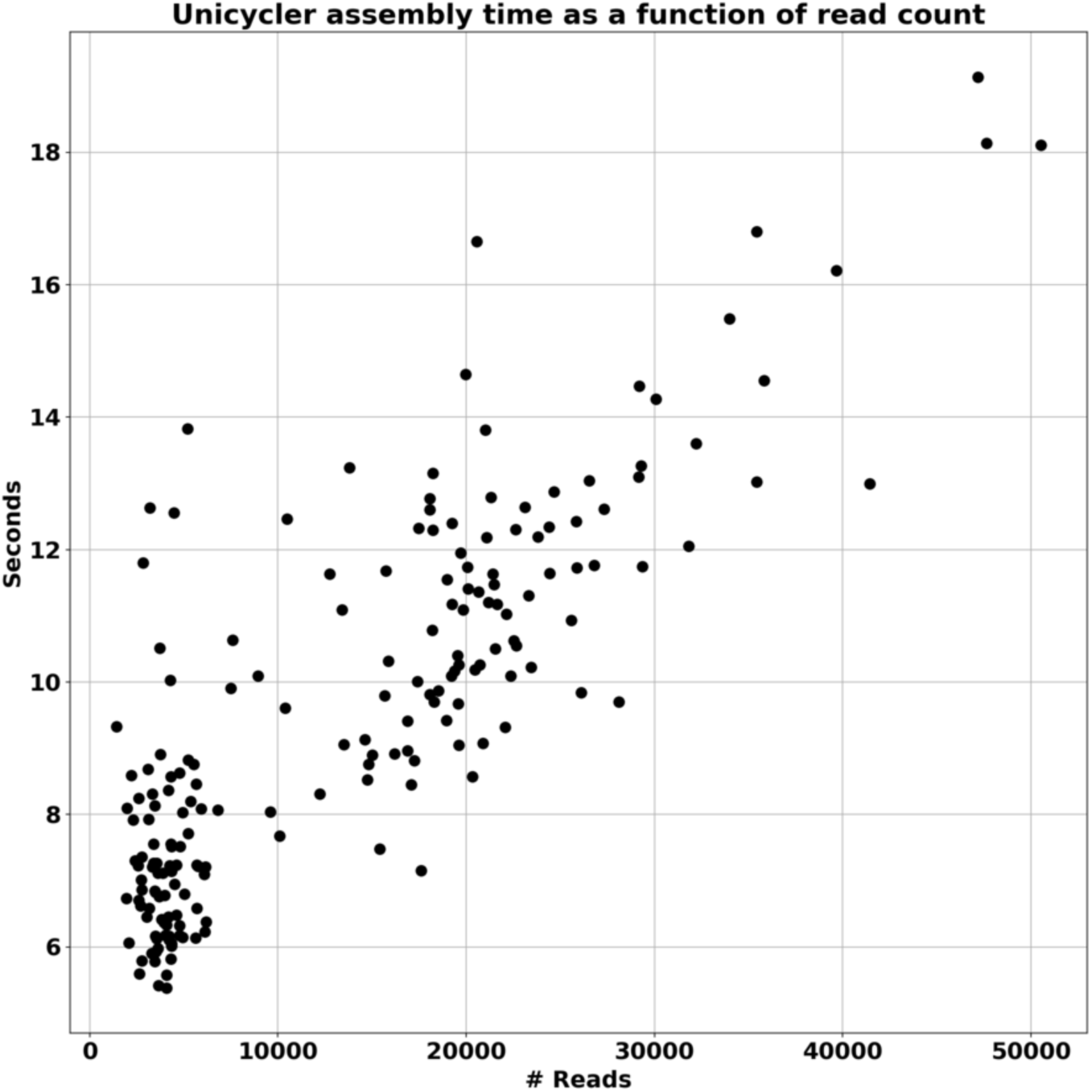
Unicycler assembly time as a function of read count. Experiments using up to 50,000 input reads show that Unicycler runtime increases approximately linearly with the number of input reads.

For 95 of the 96 experimental samples, our pipeline’s assembled contig matched the sequence generated by seqWell using their proprietary plasmid assembly workflow. For the one exception, seqWell was unable to provide an assembly, reporting that it was due to poor read quality. In this case, our pipeline was able to generate an assembly successfully after the quality filtering step eliminated more than 95% of the total read pairs.

72 of our 96 samples are clones derived from plasmids that were previously sequence-verified by Twist. It is worth noting that 48 of these 72 plasmids had the capacity to form substantial hairpins resulting in practically uninterpretable Sanger sequencing reads. For these previously sequence-verified samples, our pipeline correctly assembled all 72 plasmids, and succeeded in identifying all of the known SNPs that were present in 36 of these 72 samples.

The remaining 24 samples sequenced contained plasmids that were constructed from non-sequence-verified parts by Gibson assembly (20). Among the 24 samples, our pipeline detected 15 that contained unknown SNPs and/or INDELs. We validated two of the 15 variants by Sanger sequencing and confirmed that all 15 agreed with the assemblies provided by seqWell.

In conclusion, our workflow resulted in the correct assembly of all 96 samples, including the identification of several known and unknown variants, even when overall read quality was very low.

### Assessing assembly quality without a reference and detecting contamination

In the previous section, we described developing and validating our pipeline by comparing the resulting assemblies to a known reference sequence. However, we built our pipeline using *de novo* assembly, so that it can also be used to assemble plasmid sequences when a reference is not available. One reference-agnostic way to assess an assembly’s potential fidelity is to map the input reads to the assembly. To determine if assembly quality could be reliably assessed in this manner, we challenged our assemblies by artificially spiking in “contaminant” reads prior to assembly. We then compared the percent match between the assembled contig and the input reads for contaminated and uncontaminated reads to establish a baseline percent match for assessing assembly quality.

We ran this procedure using different proportions of contaminant reads, from 50% down to 5%, and evaluated 500 assemblies at each proportion. Each contamination simulation experiment yielded four possible outcomes (Figure 4a): (1) the assembly matches the primary reference, (2) the assembly matches the contaminant reference, (3) the assembly matches neither, or (4) the pipeline yields a fragmented assembly containing more than one contig. By varying the relative proportions of primary and contaminant reads in the combined data sets, we assess the impact of SNPs, INDELs, and other forms of contamination and evaluate the assembly pipeline’s sensitivity to contaminated data as shown in Table 1.

**Table 1:** Contamination Test Summary. Each row of this table represents the average result from a single simulation experiment (repeated 500 times). Each experiment differed by the amount and type of contamination simulated. Variant Type denotes by what metric the “contaminating” sequence (reads from a different sample that were artificially spiked) differs from the “primary” sequence (the reference sample). %Contamination refers to the percentage of the read library derived from contaminating reads. %Correct denotes the percentage of correct assemblies yielded. %Fragmented denotes the percentage of fragmented assemblies yielded. %Mismatches denotes the percentage of assemblies that did not exactly match the reference.

| Variant Type | %Contamination | %Correct | %Fragmented | %Mismatches |
| --- | --- | --- | --- | --- |
| none | 0 | 100% | 0% | 12.29% |
| 1 SNP | 20% | 100% | 0% | 13.38% |
|  | 35% | 96% | 0% | 14.41% |
|  | 50% | 44% | 0% | 14.68% |
| 2 SNPs | 20% | 100% | 0% | 14.35% |
|  | 35% | 96% | 0% | 14.82% |
|  | 50% | 32% | 0% | 15.96% |
| 3 SNPs | 20% | 100% | 0% | 14.04% |
|  | 35% | 96% | 0% | 15.21% |
|  | 50% | 35% | 0% | 16.58% |
| 4 SNPs | 20% | 100% | 0% | 14.85% |
|  | 35% | 90% | 0% | 16.83% |
|  | 50% | 18% | 0% | 17.99% |
| 608bp indel | 10% | 99% | 1% | 13.49% |
|  | 20% | 89% | 11% | 14.78% |
|  | 35% | 16% | 84% | 17.28% |
|  | 50% | 2% | 95% | 21.00% |
| unrelated | 10% | 86% | 4% | 18.24% |
|  | 20% | 28% | 72% | 23.96% |
|  | 35% | 0% | 100% | NA |
|  | 50% | 0% | 100% | NA |

**Figure 4.**
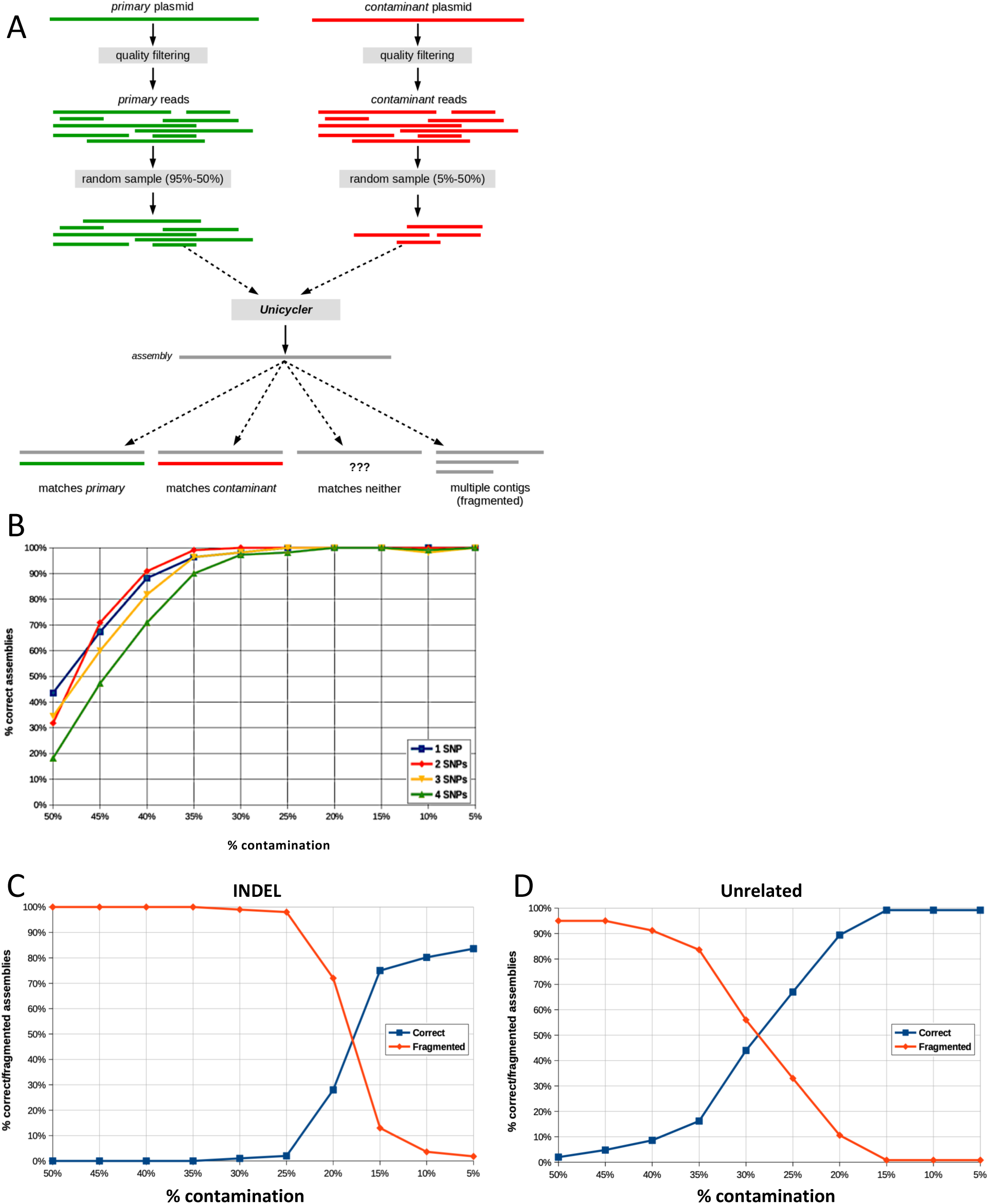
Determining the reliability of assemblies in the presence of contaminating reads. **(A)** Workflow for creating a set of reads (“primary”) with a specific proportion of contaminating reads to simulate how the pipeline is affected by contamination. Datasets of reads with 5-50% contaminating reads were created, assembled, and verified against the reference sequence. **(B)** Percentage of correct assemblies when contaminating reads contained 1-4 SNPs. Percentage of correct and fragmented assemblies when **(C)** a 608 bp INDEL was introduced into contaminating reads, or **(D)** contaminating read sequences are completely unrelated to primary sequence.

In Table 1 we report the average proportions of correct, fragmented and mismatching assemblies. Not surprisingly, as the proportion of contaminant reads increases, the proportion of assemblies correctly matching the primary reference sequences decreases. When the difference between the contaminant differs from the primary plasmid by one to four SNPs, fully 90% of assemblies match the correct primary sequence at 35% contamination or less (Table 1, Figure 4B).

When the contaminating sequence contains an INDEL, we observe a sizeable increase in the number of fragmented assemblies (Table 1). It is worth noting that in some cases the fragmented assembly consists of just two sequences: the correct sequence for the smaller of the primary and contaminant source sequences, and a second sequence that matches the insertion. When the primary and contaminant sequences differed by a 608 bp INDEL, the pipeline failed to yield correct assemblies consistently when over 20% of reads were contaminating (Table 1, Figure 4C).

When the primary and contaminant populations come from plasmids with very little sequence similarity, the resulting assemblies are more severely affected than with SNPs or INDELs. As with INDELs, the high assembly fidelity was achieved only when contamination levels were below 20% of total reads (Figure 4D). At that ratio, the assembly may be correct up to 86% of the time (Table 1). Similarly, we find that fragmented assemblies nearly disappear.

These results suggest that our pipeline produces robust results when contaminating reads are below 20% to 35%, depending on the type of contaminant and its similarity to the primary sequence. However, even when the contamination is limited enough to yield exactly one contig, it is still important for the user to know whether any contamination is present in the sample. To predict contamination likelihood in a reference agnostic way, we mapped the input reads back to the assembly for contaminated and uncontaminated samples. We predicted that, because contaminating reads will not match, this approach would provide a potential measure of contamination likelihood.

We first note that, even after filtering, some reads may not map exactly even to a perfect assembly derived from an uncontaminated sample (33-35). However, the mismatch percentages were, as we predicted, higher for read data with contamination (Figure 5A, above).

**Figure 5.**
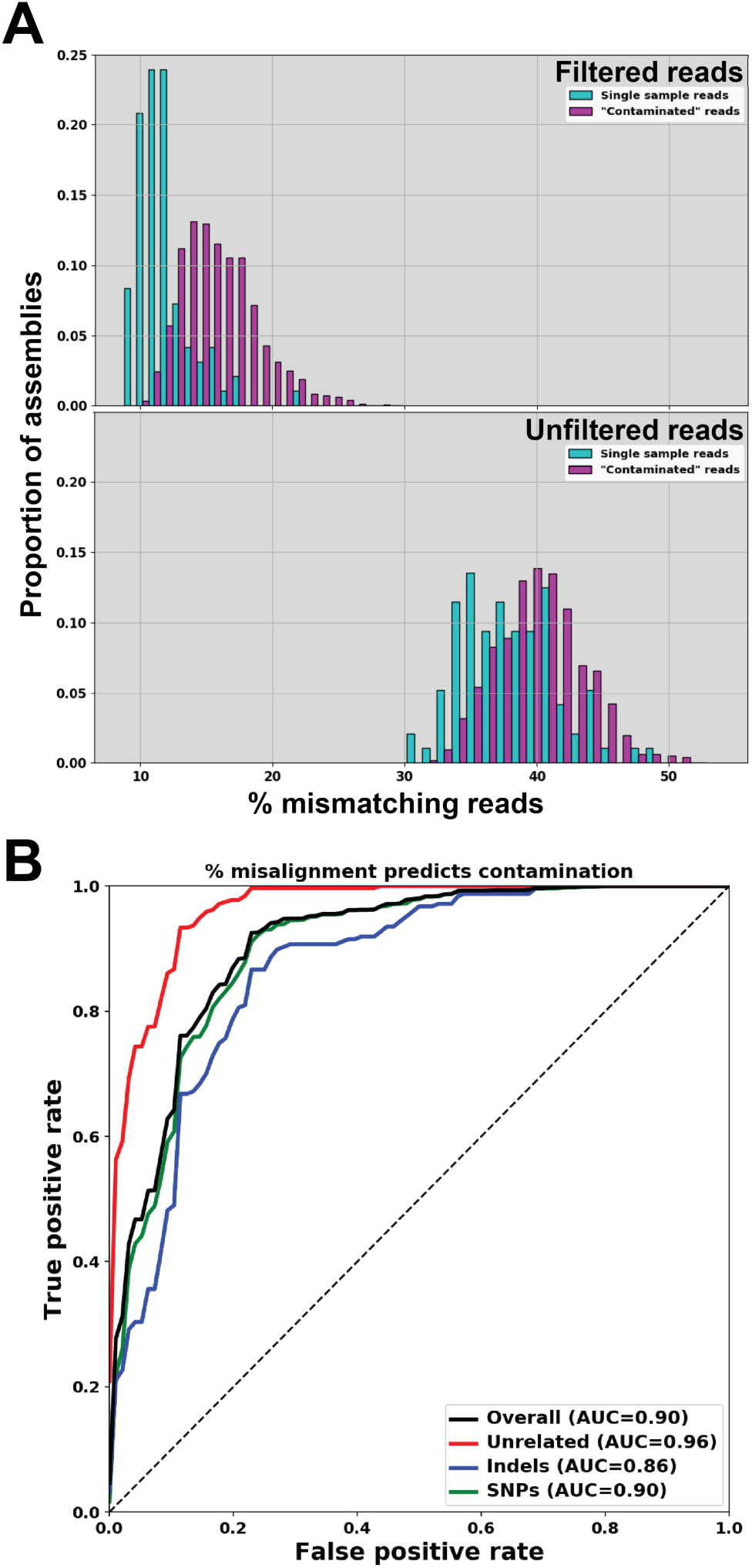
Misalignment of reads to the assembly can be used as a marker of contamination. **(A)** Input reads aligned back to their associated *de novo* assembly for contaminated and clean samples for filtered (top) and unfiltered (bottom) reads. **(B)** Receiver operating characteristic (ROC) curve demonstrating that, using the percent of mismatching reads alone, we can predict contamination with high accuracy. Here, true positives are assemblies correctly predicted as contaminated, and false positives are clean assemblies incorrectly predicted as contaminated. AUC: Area under the curve.

For the 96 assemblies described above, we find that on average 11.7% of filtered input reads fail to map exactly to their associated assembly (Figure 5A, above). With contaminated data we find that overall 15.9% of input reads fail to map to an assembly. Interestingly, without the filtering step, the same percent of contaminated and uncontaminated reads mapped to the assembly (Figure 5A, below), suggesting that a filtering step is needed to detect contamination. Further, we find that the percentage of misaligned reads is highly predictive of contaminated data, even when the contamination is relatively subtle, as with SNPs (AUC 0.86-0.96, Figure 5B). The percent of mismatching reads to the assembly thus We use this metric in our pipeline as a measure of the quality of the assembly and the likelihood of contamination.

## Discussion

We have demonstrated the feasibility of rapidly producing highly accurate plasmid assemblies from short read sequencing data by *de novo* assembly. Our publicly available workflow relies on Unicycler (27), an open source tool that was developed for assembly of circular genomes. We identified Unicycler parameters and determined read filtering thresholds that are optimal for plasmid assemblies.

We incorporated into our pipeline a novel method for detecting contamination by analyzing the percent of input reads that map exactly to locations in the final assembly. In general, our pipeline can generate accurate assemblies with fewer than 1000 reads, regardless of quality; but if we focus on high-quality reads, we can begin to detect contamination. We find that only a small fraction of high-quality reads fail to map to an assembly, while increased levels of non-mapping reads correlate with contamination. By applying strict filtering, we reduce the errant reads. We find that the resulting percent-matching metric can provide coarse information about whether the input data are likely to be contaminated. Despite our method’s simplicity, our experiments show that it is surprisingly adept at discriminating between “clean” and contaminated assemblies, demonstrating sensitivity even to contaminating plasmids with single-nucleotide mutations at relatively low levels.

The metric we use to detect contamination in single-contig assemblies is a rudimentary first step. We envision future improvements in several directions for this work. First, our rudimentary metric could be replaced with a more sophisticated machine learning approach that incorporates features of mapped and unmapped reads, such as the number of mismatching bases or their associated quality scores. Second, either this machine learning model or our realignment method could aid in selecting a contig that matches the correct plasmid sequence in cases where Unicycler produces more than one contig. Third, we wish to find ways for our pipeline to provide greater detail regarding the likely nature of any contamination it detects. We believe our likelihood score will be useful, but reporting the nature of any contamination detected would make it easier for users to assess their samples more fully.

These results have important implications for basic and applied research. Contamination and sample misidentification are rampant problems in mammalian cell research (36-39). While comparably less attention has been paid to the potential for similar problems in model organisms and *in vitro*, it has been demonstrated that many plasmids currently in circulation do not match their supposed references (40, 41). To ameliorate this problem, researchers can sequence their plasmids relatively inexpensively by coupling them to other sequencing runs at a local core facility and then using our pipeline to ensure that plasmid sequences are correct before running any experiments. seqWell has made the chemistry they use for plasmid sequencing available as library prep kits (∼$16/sample, https://seqwell.com/products/), which should facilitate the process of plasmid verification by Illumina sequencing. This small up-front cost could save considerable time and resources, as well as limiting reproducibility issues.

The ability to rapidly and automatically validate plasmids by Illumina sequencing will be especially applicable to large-scale plasmid production projects associated with synthetic biology (42). The rapidly decreasing cost of DNA synthesis and sophisticated computer-aided design tools (43, 44) have facilitated disciplined factorial experiments involving large libraries of plasmids with various genetic parts (45) for applications such as improved genetic circuit design (46), multiplexed CRISPR-Cas (47) or metabolic engineering (48). These workflows, which are often automated (49, 50), necessitate an automated sequence verification pipeline like the one we’ve described.

Further, this pipeline will be an asset to another large-scale application: high-throughput plasmid libraries. Increasingly, high-throughput screens are being performed from libraries of hundreds to thousands of genes encoded on plasmids or lentiviral “transfer plasmid” genomes. Panels of this sort are so robust that they have made it possible to specify each gene involved in a molecular pathway (51-53) in a single screen, work that would have previously taken years to decades. Notably, plasmid libraries have been shown to perform sgRNA CRISPR/Cas9 knock-out (51) or shRNA knock-down (54) screening panels (Reviewed in (55)).

Though these technologies were introduced in 2014 and 2011, respectively, they have been slow to be adopted, in part we believe from the difficulty of generating plasmid libraries. Though, the bottleneck of these types of screens is still generating a plasmid library, which includes the essential step of verifying plasmid sequences. The plasmid verification pipeline described here can solve a critical need to make generating high throughput screening libraries faster, cheaper, and therefore more feasible.

This work is also relevant for data security (56, 57). There is increasing interest in using DNA as a medium for information storage (58, 59). Recently, researchers began experimenting with storing information in plasmids, as they can be rapidly replicated (60, 61). A technology for encrypting and securing digital information using cybersecurity techniques has also been developed for plasmids (62). Maintaining digital references for these sequences would defeat the purpose of storing the information in DNA, so these efforts will be greatly facilitated by the rapid *de novo* assembly pipeline we’ve described.

The described pipeline was built using open source tools and is available to the community through the supplement.

## Supplemental Information

The bioinformatics pipeline is licensed open-source can be found here: https://bitbucket.org/genofabinc/oss/src/master/denovo/

## Acknowledgements

This work was supported by NSF Awards 1934573 and 1832320. The funder had no role in study design, data collection and analysis, decision to publish, or preparation of the manuscript.

